# Loss of protein stability and function caused by P228L variation in NADPH-cytochrome P450 reductase linked to lower testosterone levels

**DOI:** 10.1101/2022.08.16.504152

**Authors:** Maria Natalia Rojas Velazquez, Mathias Noebauer, Amit V. Pandey

## Abstract

Cytochrome P450 oxidoreductase (POR) is the redox partner of steroid and drug-metabolizing cytochromes P450 located in the endoplasmic reticulum. Mutations in POR cause a broad range of metabolic disorders. The POR variant rs17853284 (P228L) identified by genome sequencing has been linked to lower testosterone levels and reduced P450 activities. We expressed POR wild type and the P228L variant in bacteria, purified the proteins, and performed protein stability and catalytic functional studies. Variant P228L affected the stability of the protein as evidenced by lower unfolding temperatures and higher sensitivity to urea denaturation. A significant reduction of model electron acceptors was observed with POR P228L while activities of CYP3A4 were reduced by 25%, and activities of CYP3A5, and CYP2C9 were reduced by more than 40% compared to WT POR. The 17,20 lyase activity of CYP17A1 responsible for production of main androgen precursor dehydroepiandrosterone, was reduced to 27% of WT in presence of P228L variant of POR. Based on *in silico* and *in vitro* studies we predict that the change of proline to leucine may change the rigidity of the protein, causing conformational changes in POR, leading to altered electron transfer to redox partners. A single amino acid change can affect protein stability and cause a severe reduction in POR activity. Molecular characterization of individual POR mutations is crucial for a better understanding of the impact on different redox partners of POR.

## 1. Introduction

Cytochrome P450 oxidoreductase (POR) (OMIM: *124015, HNGC:9208) has a key role in several metabolic processes [1, 2]. POR is located in the endoplasmic reticulum of the cells, where it operates as an electron donor for the cytochromes P450, heme-oxygenase, squalene mono-oxygenase, and other redox partners [2-4]. POR has distinct domains that bind cofactors flavin adenine dinucleotide (FAD) and flavin mononucleotide (FMN) which are connected by a flexible hinge region. POR catalyzes the transfer of electrons from the nicotinamide adenine dinucleotide phosphate (NADPH) through FAD to FMN and then to its redox partners or substrates. Microsomal cytochromes P450 perform a range of metabolic reactions including drug metabolism, biosynthesis of steroid hormones, biosynthesis of sterols, and metabolism of retinoic acid, all of which require POR for their catalytic function. A disruption of POR activity affects the catalytic function of several enzymes in steroid production including the 21-hydroxylase (CYP21A2) for mineralocorticoids and glucocorticoids production, the 17-hydroxylase (CYP17A1) for the synthesis of androgens, and the aromatase (CYP19A1) responsible for estrogen production (**Figure 1**) [5-7].

**Figure 1.**
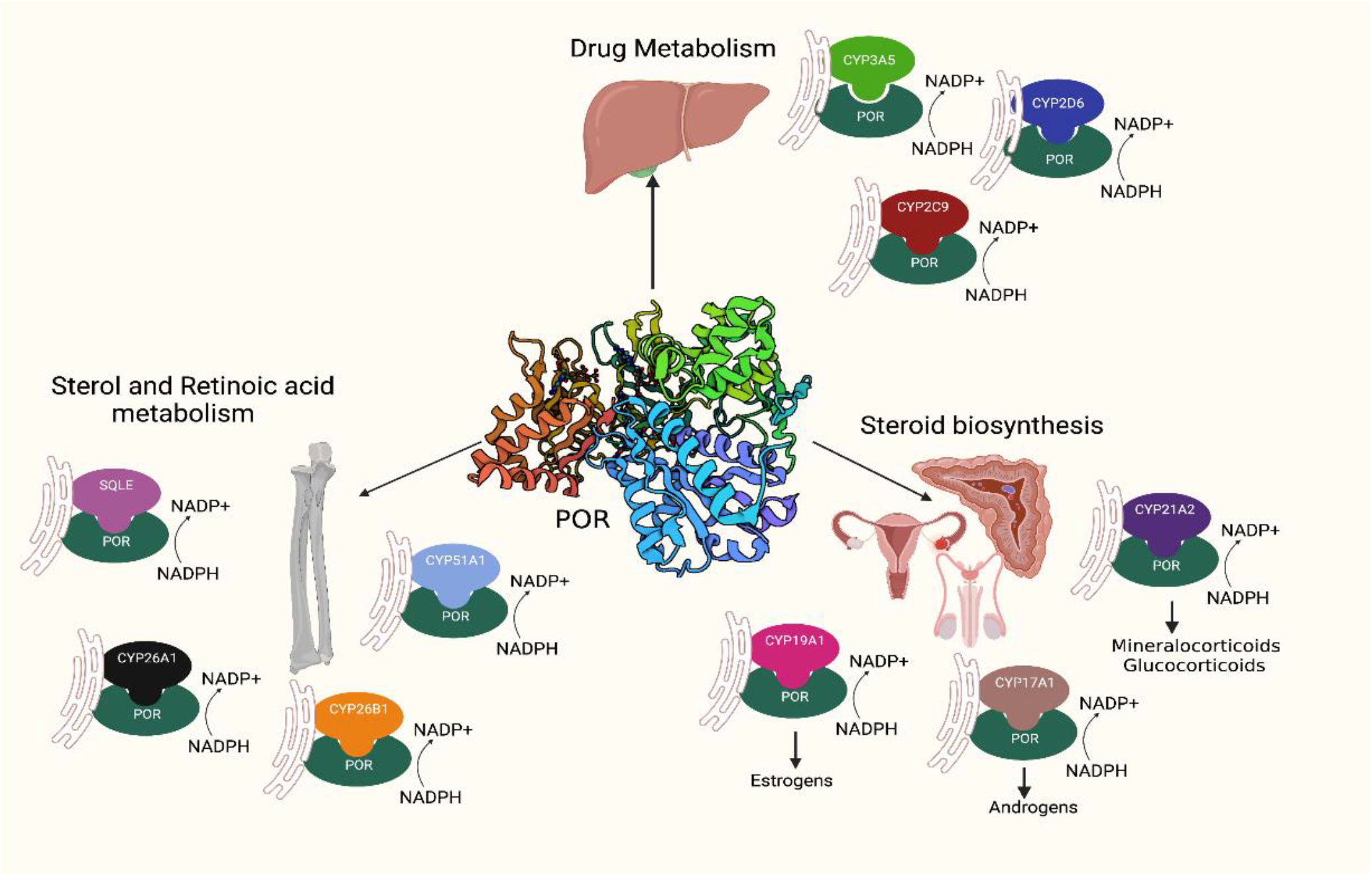
POR is a key-enzyme in several metabolic processes, such as drug metabolism, steroid biosynthesis and sterol and Retinoic acid metabolism. POR provides the necessary equivalents from NADPH to its multiple redox partners to accomplish their functions.

POR is expressed in all tissues, showing ubiquitous expression in the liver, adrenal glands, gonads, and others [8]. Human POR consists of 680 amino acids and is a membrane-bound protein coded by the *POR* gene located in chromosome 7 with 15 coding exons and 1 non-coding exon. Mutations in POR can lead to Cytochrome P450 oxidoreductase deficiency (PORD; OMIM 613571 and 201750), a rare form of congenital adrenal hyperplasia, disrupting the steroid production in the adrenal and gonads [2, 9]. This disruption may lead to genital ambiguity at birth in both males and females [9, 10]. Patients with PORD may develop skeletal deformities resembling Antley-Bixler Syndrome [4, 5, 10-12]. Additionally, several milder symptoms have been described in patients with PORD due to the multiplicity of its redox partners [5].

After the first cases of PORD were described in 2004, a large number of POR variants have been reported [1, 13-15]. From the data available from big genome projects, hundreds of single-point mutations have been identified not only in patients but also in non-clinical populations [1, 4, 5, 16]. Henceforth, our group started to study how these common variants could affect the healthy carriers without an evident phenotype, especially at the level of drug metabolism. Previous studies of our group have shown that polymorphisms in POR can alter the enzymatic activities of its redox partners [6, 17-21].

In the current study we have performed enzymatic and structural analysis of the rs17853284 (NM_000941.3(POR):c.683C>T, POR P228L) variant of POR found in apparently healthy individuals in big genomic studies such us 1000 Genomes, BioMe Biobank, Genotype-Tissue Expression Project (GTEx), and Human Genome Diversity Project. Recent genome-wide association studies have indicated a link to lower testosterone levels with the presence of POR variant rs17853284 [22, 23]. We performed bioinformatics studies to predict the impact of the mutation on protein stability and function and produced recombinant WT and P228L variants of POR in bacteria, purified the proteins using immobilized-metal affinity chromatography (IMAC) and performed enzyme kinetic experiments with small molecules and cytochrome P450 enzymes together with protein stability assays. Our results show lower activities of several redox partners with the POR variant P228L, providing confirmation for its links to lower testosterone levels and indicating that together with a severe variant on another allele, the POR rs17853284 may result in POR deficiency.

## 2. Results

### 2.1 Genetic distribution of POR P228L

We identified the POR variant P228L as an interesting target for further studies by an extensive search in the genome databases where there is a conflicting interpretation of its pathogenicity. Previous studies have shown that P228L affects the activity of some of its redox partners [1, 5]. The P228L POR is found as a rare variant in the NCBI database. The allele frequency varies between T= 0.00104 to T=0.00346 among the different databases (around 0.2 % of the total number of alleles sequenced) (**Table 1)**. For the population distribution of P228L, we extracted the data from GnomAD database. The P228L (variant ID: 7-75610876-C-T) appears to be a rare variant mainly identified in the European population (allele frequency= 0.004702), while it is found to a lesser degree in African/African Americans (allele frequency= 0.001057) and Latino/Admixed Americans (allele frequency= 0.0007576). The allele frequency decreased to 0.0001282 in South Asians and the P228L variant of POR has not been found in East Asians (**Figure 2)**. In two recent genome-wide association studies a link to lower testosterone levels has been associated with the POR variant rs17853284 [22, 23].

**Table 1.**
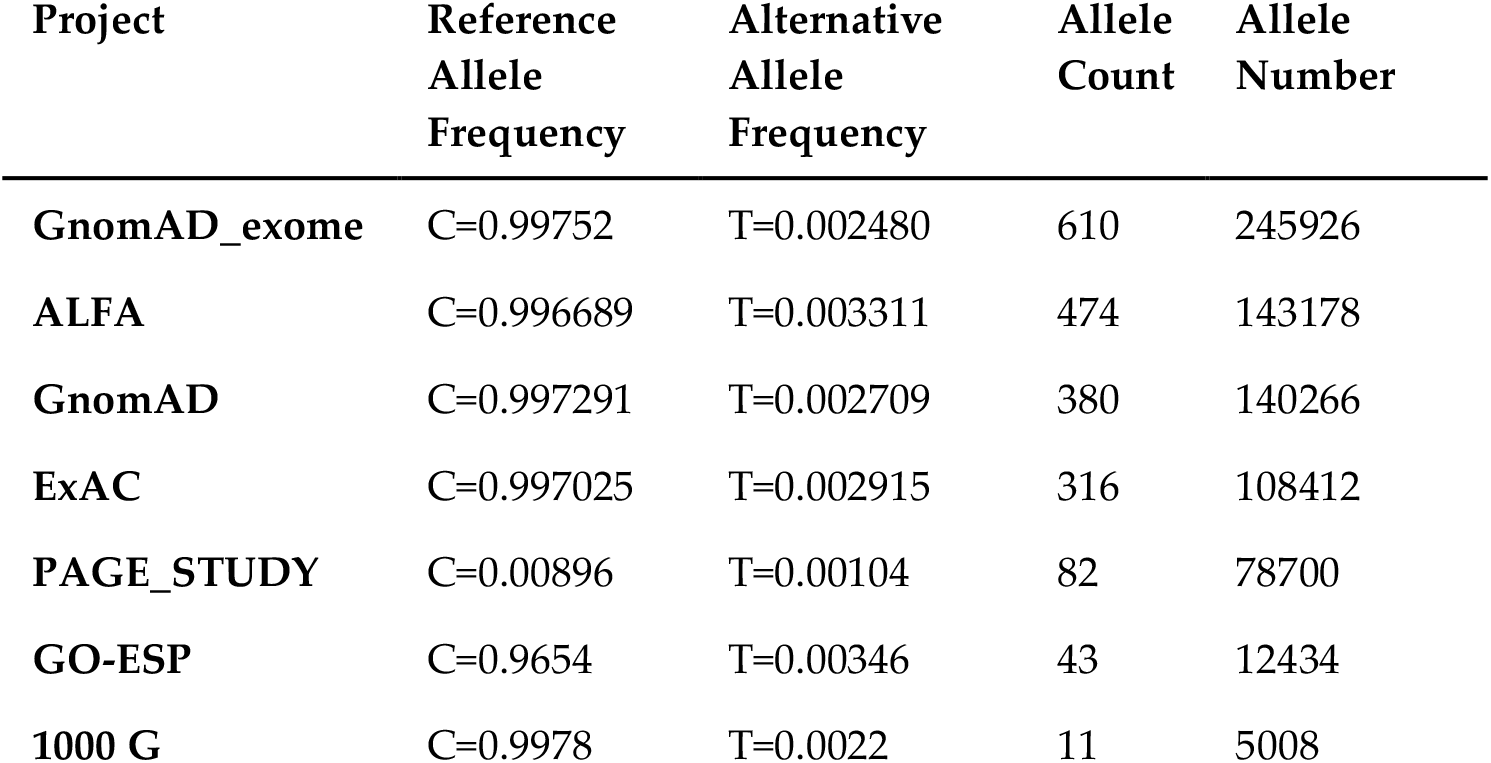
Allele frequencies of POR P228L (DBSNP ID: rs17853284) from different genome projects. Data obtained from NCBI database.

**Figure 2.**
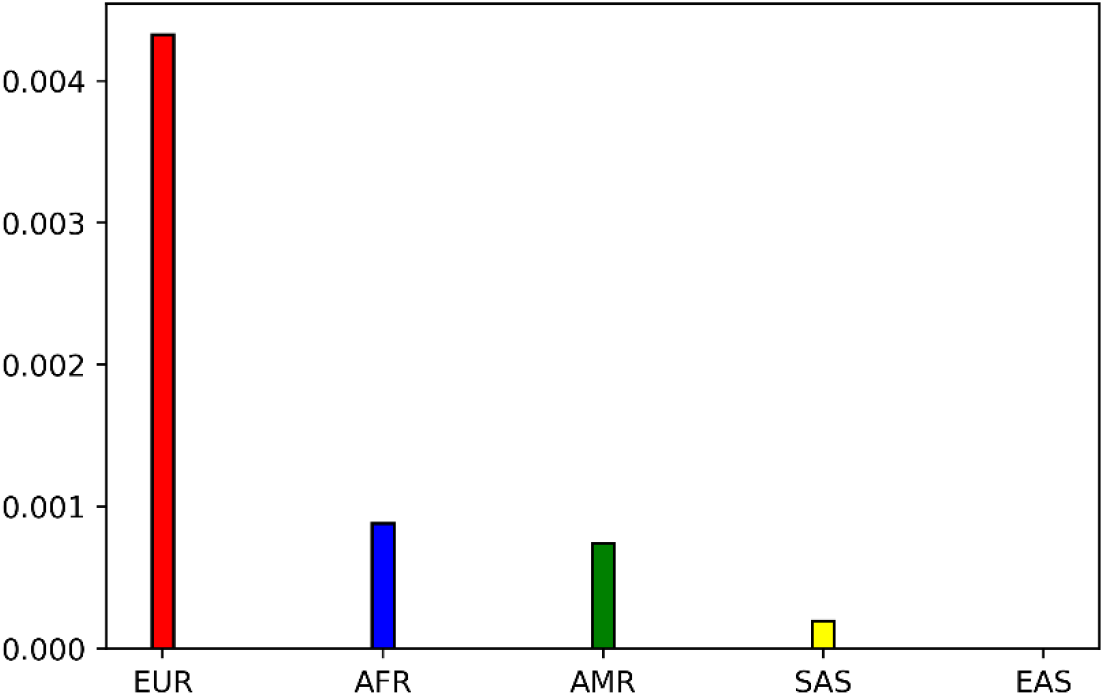
Population distribution of POR variant P228L extracted from GnomAD database (variant ID: 7-75610876-C-T). P228L appears to be a rare variant mainly found in Europeans (EUR). The allele frequency is similar in African/African Americans (AFR) and Latino/Admixed Americans (AMR). The frequency is lower in South Asians and P228L has not been found in East Asians (EAS).

### 2.2 Conservation and structural analysis of POR

We assessed the sequence and structure conservation of POR at position P228 by ConSurf analysis using 250 homologous sequences of POR across species. The position P228 has a score of 9 being classified as highly conserved (**Figure 3)**. The P228 is located at the surface of the protein in a highly conserved region close to the FMN domain and is predicted to affect the flexibility of the POR molecule which is essential for the transfer of electrons and interaction with redox partners. Usually, conservation across species indicates an important role of the amino acid for the function of the protein [1, 5, 24].

**Figure 3.**
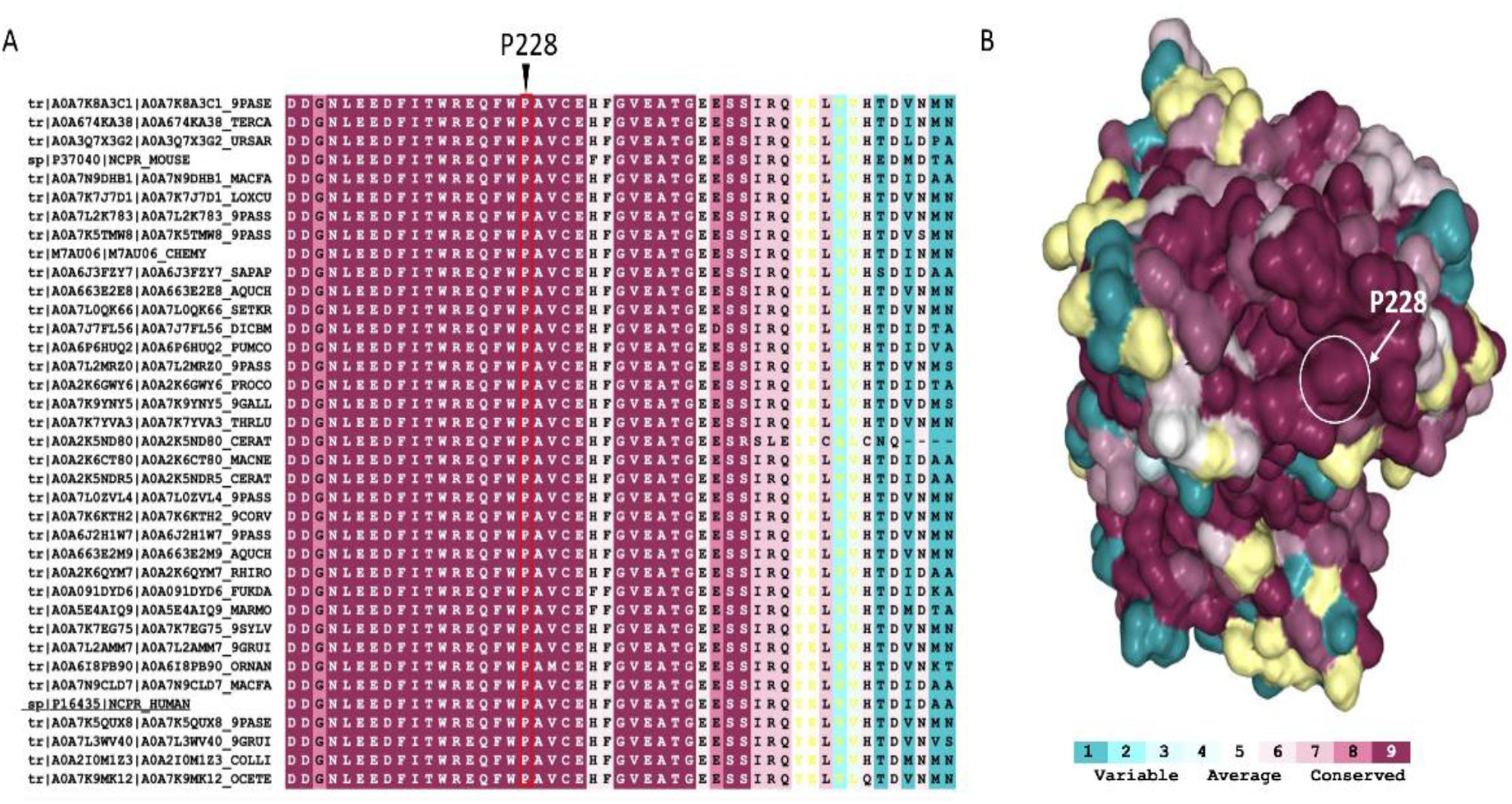
ConSurf analysis of POR (PDB # 5FA6) evolutionary conservation profile. **(A)** A Segment (residues 210-258) of the multiple-sequence alignment of 250 POR homologues across species. Position 228 is indicated with an arrow; the P228 has a score of 9 being classified as highly conserved. The alignment is colored according to the ConSurf conservation color bar (details below). **(B)** POR structure is illustrated as a surface model; the amino acids are colored according to the conservation scores. The color bar varies from (1) cyan meaning variable to (9) maroon meaning conserved. The amino-acids colored yellow has not been classified due to insufficient data.

### 2.3 In silico predictions of POR P228L stability

We checked the possible impact of the mutation P228L on the functionality of POR using seven sequence-based tools. As shown in **Table 2**, the P228L variant of POR is predicted to be deleterious in all of them since these are mostly based on sequence conservation, and therefore, are in agreement with the ConSurf analysis. Additionally, we used DynaMut-2.0 to predict the vibrational Entropy difference (ΔΔS) which was found to be -0.904 kcal.mol^-1^K^-1^ between POR wild-type and the P228L variant calculated with [ENCoM](https://github.com/NRGlab/ENCoM) (**Figure 4)**. A subtle destabilizing effect on POR structure was predicted (predicted ΔΔG -0.64 kcal/mol) due to P228L variation causing decreased flexibility of the molecule. With this information, we can conclude that P228L POR may undergo conformational changes which might decrease its function by affecting the transition between the open and closed conformations of POR. A closed conformation of POR, when the FAD and FMN domains are near each other, is required for the electron transfer from FAD to FMN [12, 25]. The open conformation of POR is needed for the binding of redox partners for the transfer of electrons [25, 26].

**Table 2.** *In silico* predictions of protein stability upon single amino acid change using sequence-based tools. Reference Human POR sequence NCBI: NP_000932.3

| Method | Prediction | Accuracy |
| --- | --- | --- |
| PredicSNP2 | Deleterious | 0.89 |
| CADD | Deleterious | 0.52 |
| DANN | Deleterious | 0.89 |
| FATHMM | Deleterious | 0.67 |
| FunSeq2 | Deleterious | 0.61 |
| GWAVA | Deleterious | 0.47 |
| Meta-SNP | Deleterious | 0.6 |

**Figure 4.**
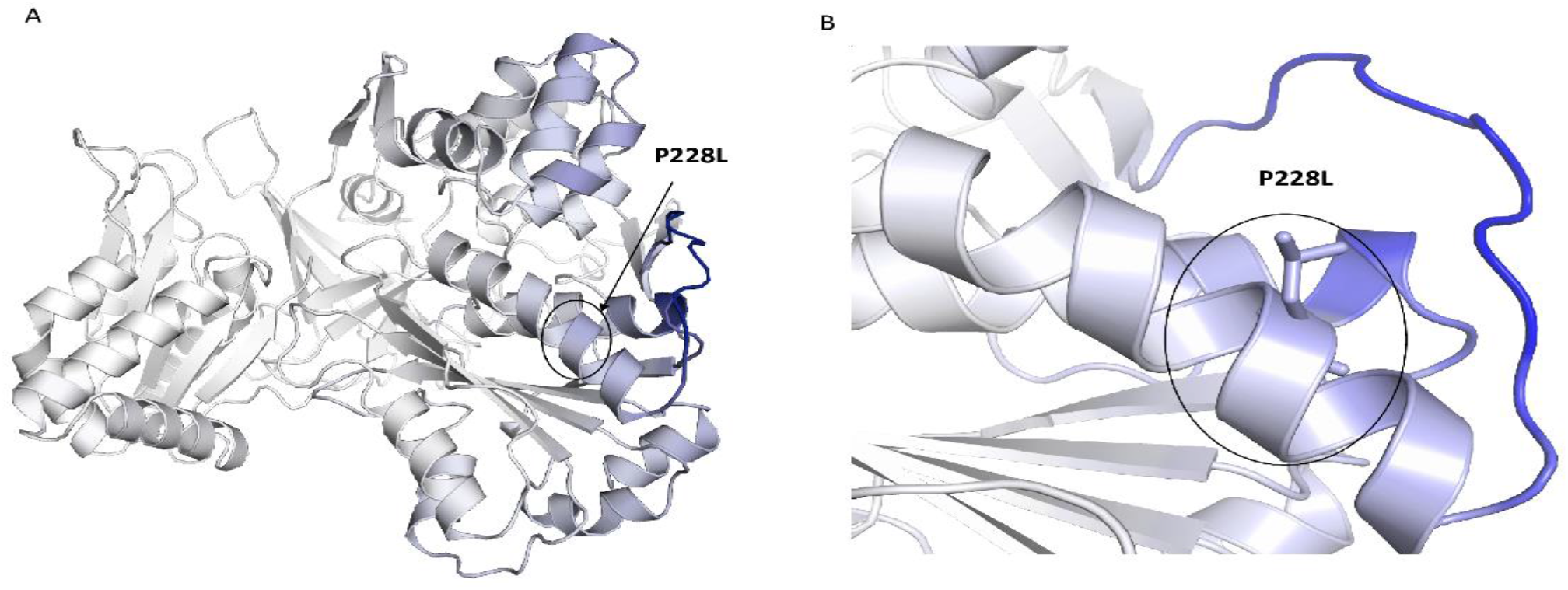
Visual representation of the Vibrational Entropy difference (ΔΔS) between POR wild-type and P228L structures (ΔΔS_Vib_: -0.904 kcal.mol^-1^. K^-1^) calculated with [ENCoM](https://github.com/NRGlab/ENCoM). Mutant P228L decreased the flexibility of the molecule. Amino acids are colored according to the vibrational entropy change upon mutation. Blue represents a rigidification of the structure and Red a gain in flexibility.

### 2.4 Flavin content of WT and mutant P228L proteins

Flavins are crucial for electron transfer within POR, and its function can be severely affected due to reduced binding of cofactors FMN and FAD [27, 28]. To test the changes in cofactor binding, the flavin contents of WT and P228L POR were measured. Flavin content of P228L (FMN 79 %, FAD 82 % of WT flavin content) was moderately lower using urea-mediated denaturation (**Figure 5, A, B**) suggesting that structural changes in the P228L variant may slightly affect the binding of FMN/FAD.

**Figure 5.**
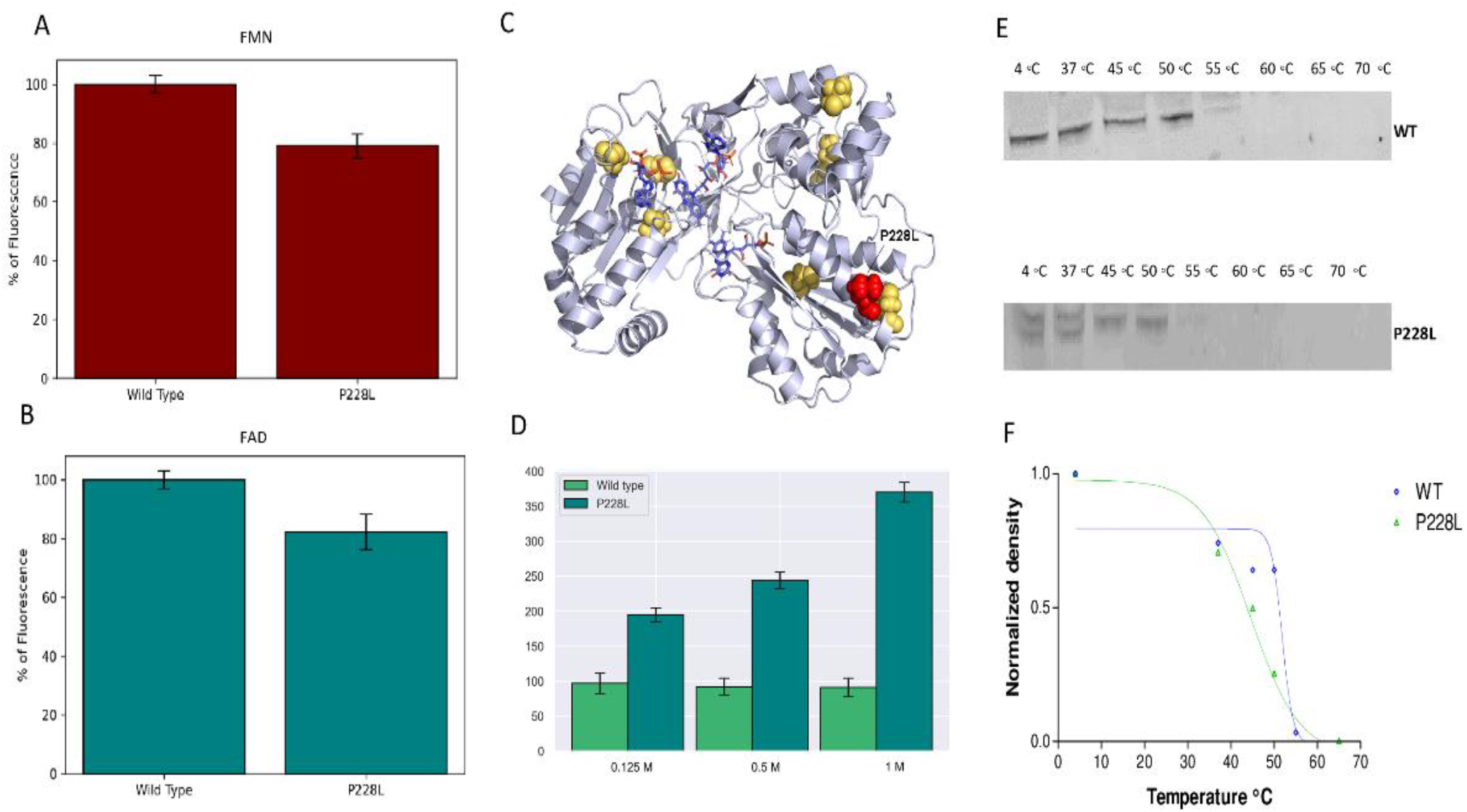
Flavin content and stability of WT and P228L POR. The fluorescence of released FMN/FAD was measured with excitation at 450 nm and emission at 535 nm (A) FMN content and (B) FAD content of POR WT and P228L are shown. (C) POR structure is illustrated showing, the cysteine residues inside or on the surface of the folded protein colored in yellow (D) Stability analysis of WT and P228L by CPM fluorescence assay. The stability of protein structure was tested by exposure to increasing concentration of Urea, the fluorescence emission at 463 nm increased upon unfolding due to increased access of CPM molecules to hidden cysteine residues in the unfolded protein. The P228L mutant of POR was less stable, the P228L unfolded at a lower concentration of urea and the flavin content was lower than WT POR.(E) Analysis of WT and P228L stability by FASTpp assay. FASTpp of WT and P228L were analyzed by Western blot as described in the methods. (F) Representative melting curve of WT (Tm 51.7◦C) and P228L (Tm 44.4◦C). The P228L mutant of POR was less stable than the WT POR protein and was degraded faster and at lower temperatures upon thermolysin treatment

### 2.5 Stability of mutant P228L proteins

The *in-silico* analysis suggested an effect on protein stability due to variation P228L in POR which was corroborated by the decrease in flavin content. Therefore, we analyzed the stability of the WT and P288L variant forms of POR by performing a fast proteolysis assay. At a range of temperatures from 4°C to 70°C, both the WT and P228L forms of POR were exposed to proteolysis by thermolysin. Thermolysin is a metalloprotease that cleaves peptide bonds at the N-terminal of hydrophobic amino acids when the protein is unfolded. The unfolding temperature is often directly related to the stability of the protein structure. Mutations that alter the structure of the protein generally result in lower unfolding temperatures [29, 30].

Western blot analysis of the FASTpp reaction with the WT and P228L forms of POR (**Figure 5, C, D**) showed that for the P228L variant, proteolysis occurred even at 4°C, and the P228L POR was degraded to a higher extent compared to WT POR. The P228L variant of POR was completely degraded at around 45°C while the WT POR can still be spotted at 52°C. This indicated that the P228L variant of POR is less stable towards proteolysis. Based on these experiments we can conclude the amino acid change P228L in POR modifies the structure and consequently the unfolding properties of the POR protein making it less stable.

Furthermore, we tested the conformation and stability of the WT and P228L mutant by diethylamino-3- (4-maleimidophenyl)-4-methylcoumarin (CPM) fluorescence assay. The CPM molecules interact with cysteine residues in the proteins. Human POR has seven cysteine residues, of which three are near the surface of the protein and the rest are buried inside the structure of the folded protein and are not accessible unless the protein unfolds (**Figure 5, C**). The WT and the P228L variant of POR were treated with three different concentrations of urea to gradually unfold the protein. In the case of the P228L variant of POR, the fluorescence signal increased with the concentration of urea but the WT POR did not show any changes at the concentrations of urea used in our experiments, suggesting that the P228L variant of POR is more sensitive to urea denaturation, and may exist in a different conformation and be less stable than WT POR (**Figure 5, D) [12, 15, 31]**.

### 2.6 Effect of POR variant P228L on cytochrome c and MTT reduction activities

To determine how the variation P228L alters the catalytic activities of POR, we tested the capacity of recombinant WT POR and P228L variants to reduce its substrates (a small protein, cytochrome c, and a small molecule, MTT). The P228L variation in POR severely decreased the efficiency of cytochrome c reduction showing a loss of 74 % of its activity in contrast to WT POR (**Table 3, Figure 6, A**). In the case of the MTT reduction assay, we observed not only a severe loss of activity (a decrease of >90 %) but also a remarkably lower affinity for the substrate (**Table 3, Figure 6, B**). The decreased capacity of POR for reduction of small molecules due to variation P228L suggests that there is a reduction in electron transport from NADPH to FMN and finally to substrates, which might be due to protein instability or impact on conformation changes from close to open forms by affecting domain movements which disturb the transfer of electrons as predicted with the *in-silico* studies [25, 26].

**Table 3.** Kinetics parameters for the reactions catalyzed by recombinant WT or P228L POR. Vmax/Km was used to compare the activity of WT vs P228L, the WT activity was set at 100%

| | $V_{max}$<br>(nmol/min/mmol of<br>POR) | $K_m$ ($\mu M$) | $V_{max}/K_m$<br>(% of WT) |
| --- | --- | --- | --- |
| Cytochrome c reduction activity |  |  |  |
| WT | 633 | 6.2 | 102.1 (100) |
| P228L | 119 | 5.6 | 27 (27) |
| MTT reduction activity |  |  |  |
| WT | 519 | 42.2 | 12.3 (100) |
| P228L | 267 | 235 | 1.1 (9) |

**Figure 6.**
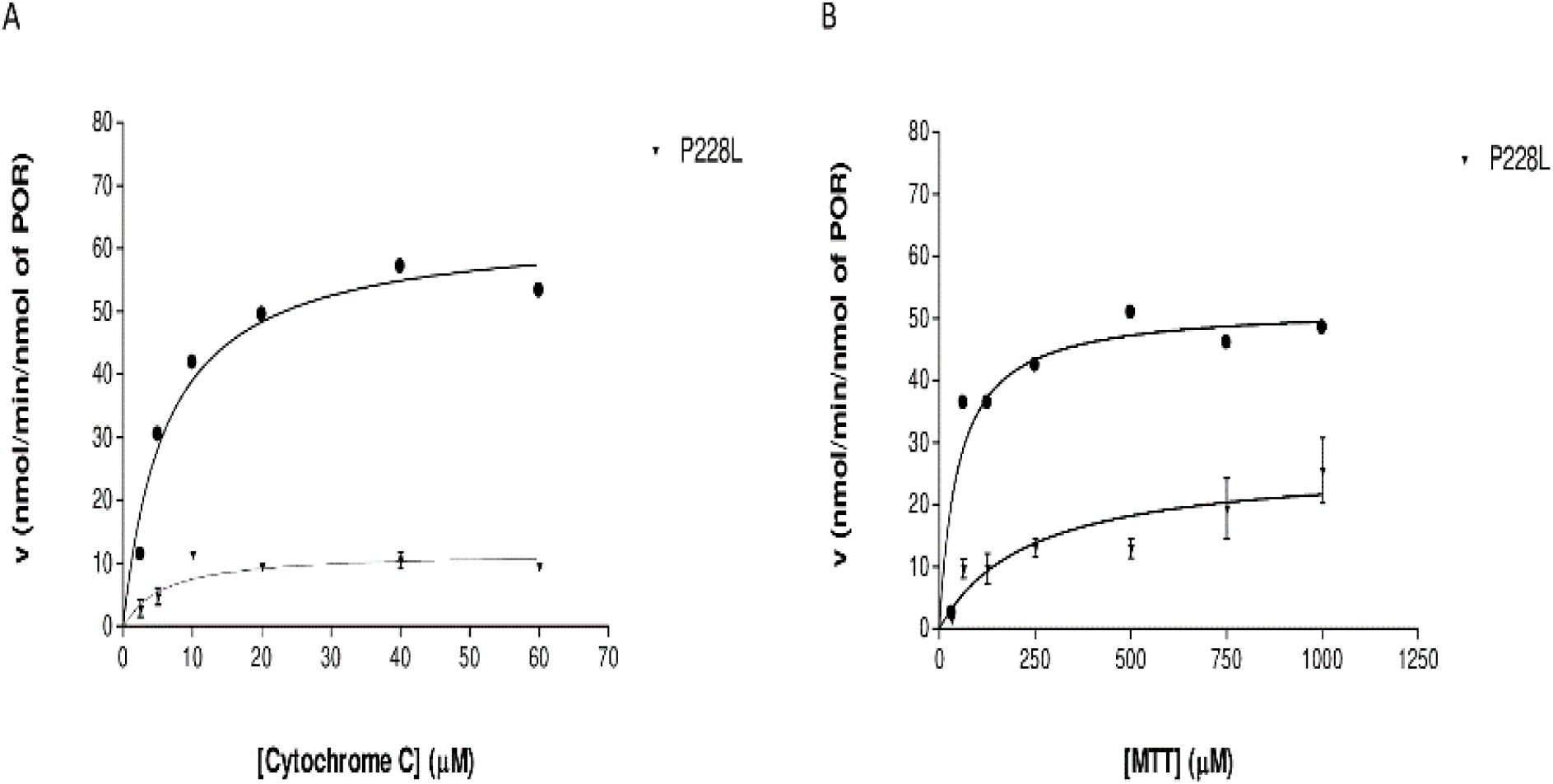
Kinetics of small molecule reduction by WT and P228L POR. (A) Cytochrome c reduction assays by WT and P228L POR. (B) MTT Reduction assay with WT and L374H variant of POR. The curves represent the best nonlinear fits to the Michaelis-Menten equation.

### 2.7 Activities of CYP3A4, CYP3A5, and CYP2C9 enzymes with POR P228L

We assessed how the P228L variation in POR affects the activity of three major drug-metabolizing enzymes CYP3A4, CYP3A5, and CYP2C9. The activity of cytochromes P450 with the WT POR was set as a hundred percent, and results are shown as a percentage of activity with WT POR. In CYP2C9 assays POR variant P228L showed 51.5 % (**Figure 7, A**) of WT activity while (B) In CYP3A5 assays variant P228L had 41 % of WT activity (**Figure 7, B**).

**Figure 7.**
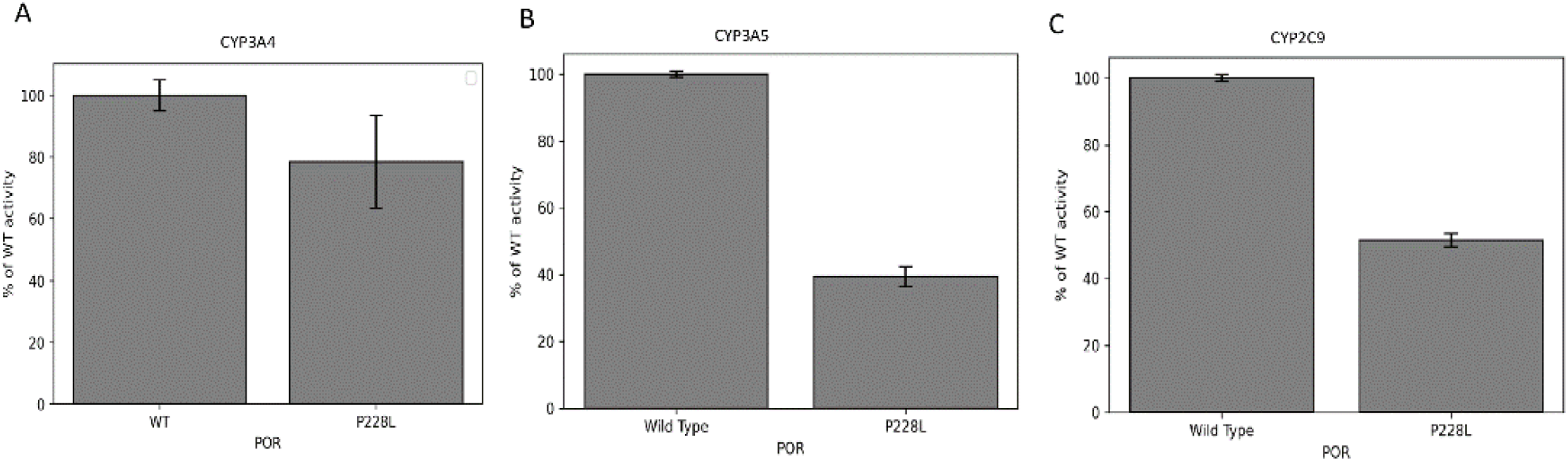
CYP3A4, CYP3A5 and CYP2C9 activity promoted by WT and P228L POR. Activity with the WT POR was set as a hundred percent, and results are shown as a percentage of WT activity. (A) In CYP3A4 assays POR variant P228L had 79% of WT activity while (B) In CYP3A5 assays variant P228L had 41% of WT activity and (C) In CYP2C9 assays POR variant P228L had 52% of WT activity.

### 2.8 Activities of steroid metabolizing CYP17A1 and CYP19A1 enzymes with POR P228L

Two recent genome-wide association studies showed a link between lower testosterone levels and the presence of POR variant rs17853284 (P228L) [22, 23]. To test the possible link between production of lower steroid precursors and testosterone level due to presence of POR P228L we checked the activities of steroid metabolizing cytochromes P450 CYP17A1 and CYP19A1. We found that conversion of progesterone to 17-hydroxyprogesterone metabolized by CYP17A1 17α-hydroxylase activity was only 57% of the WT when supported by P228L variant of POR (**Table 4**). The 17,20 lyase activity of CYP17A1 measured by conversion of 17α-hydroxy pregnenolone to dehydroepiandrosterone (DHEA) was found to be only 27% of the WT when supported by P228L variant of POR (**Table 4**). Lower amounts of androgen precursor DHEA would lead to lower substrate availability for downstream enzymes involved in androgen production, leading to overall testosterone levels, explaining the linkage seen between lower testosterone and rs17853284 variant of POR. We also measured the activity of CYP19A1 involved in estrogen production and found 54% reduction in activity in presence of P228L variant of POR (**Table 4**), suggesting estrogen production may also be affect by the rs17853284 variant of POR.

**Table 4.** Calculated kinetic parameters for activities of CYP17A1, CYP19A1 and supported by WT and P228L POR. Comparison of the steroidogenic enzyme activities supported by WT or P228L mutant of POR. All data are Vmax/Km, shown as a percentage of the wild-type control, set at 100%.

| | Vmax<br>(pmol/min*mg) | Km<br>(μM) | Vmax/Km<br>(% of WT) |
| --- | --- | --- | --- |
| ***CYP17A1: 17-hydroxylase activity (Prog to 17OHP)*** |  |  |  |
| **WT** | 257 ± 32 | 4.1±0.5 | 63 (100) |
| P228L | 193 ± 19 | 5.3±0.7 | 36 (57) |
| ***CYP17A1: lyase activity (17OHPreg to DHEA)*** |  |  |  |
| **WT** | 59.32 ± 12.35 | 1.34±0.22 | 44 (100) |
| P228L | 34.13 ± 7.09 | 2.82±0.39 | 12 (27) |
| ***CYP19A1: aromatase (D4A to E<sub>1</sub>)*** |  |  |  |
| **WT** | 12.35 ± 1.65 | 0.27±0.06 | 46 (100) |
| P228L | 7.62 ± 1.64 | 0.37±0.08 | 21 (46) |

## 3. Discussion

With the emergence of next-generation sequencing technologies, several big sequencing projects have generated large amounts of population genetic data and POR variations were not only found in patients with steroid disorders but also in healthy individuals [1, 5, 32]. Consequently, we became interested in how some of these variants could affect the metabolic functions of POR without causing an evident phenotype in the carriers. Especially, we wondered if these variations in POR could alter the metabolism of drugs since POR is required for nearly all cytochromes P450 located in the endoplasmic reticulum and is necessary for the metabolism of drugs and xenobiotics [13, 28, 33].

In this study we focused on the POR variant P228L (rs17853284) that was found in healthy individuals especially those of European origin (**Figure 2, Table 1**), mainly in one allele, or in combination with another allele (P228L + A503V) [28]. The POR P228L is classified as of conflicting interpretation for its pathogenicity since previous studies have shown the variable impact on the activity of its redox partners [34, 35]. The P228L POR showed similar activity to WT supporting heme oxygenase-1 that is involved in the catabolism of heme [34]. The CYP17A1 and CYP19A1 are key enzymes in the biosynthesis of steroids [35]. We tested and confirmed that production of androgen precursor DHEA was significantly lower in presence of POR P228L, explaining the recently found associations of lower testosterone levels linked to rs17853284 (POR P228L) variant of POR, since a general reduction in androgen precursors will create lower substrate levels for downstream enzymes involved in production of testosterone. With the cytochromes P450 located in the liver, the P228L variation in POR has shown a considerably lower catalytic capacity than WT decreasing by 80% the activity of CYP1A2 and by 60% the activity of CYP2C19. The CYP1A2 is involved in the metabolism of caffeine and other xenobiotics while CYP2C9 metabolizes about 10 % of the clinically relevant drugs [13, 32].

In the present study, the *in-silico* analysis revealed that the position P228 in POR is highly conserved across species, therefore, changes in this position could affect the structure and function of the protein (**Table 2, Figure 3-4**). In the structural studies, P228L showed a reduction of 20% in the binding of flavins (**Figure 5, A,B)**, which can be correlated with loss of protein function since both the FAD and FMN play a fundamental role in electron transfer. In addition, the P228L variant of POR appeared more susceptible to unfolding by both chemical denaturation (urea) and proteolysis (thermolysis), indicating changes in protein conformation and loss of stability (**Figure 5, C-F)**. Furthermore, the P228L variant of POR had a considerably lower capacity to reduce its small molecule substrates (Figure 6, Table 3) and a significant decrease of between 25 to 60 % of CYP3A4, CYP3A5 and CYP2C9 enzymatic activities supported by P228L variant of POR was observed compared with POR WT. Based on in silico and in vitro studies we hypothesized that the change of proline to leucine at amino acid position 228 in POR might enhance the rigidity of the protein structure hindering the conformational changes of POR, that affect electron transfer to the redox partners. A severe reduction in CYP17A1 17,20 lyase activity would lower the levels of androgen precursor DHEA potentially affecting the production of testosterone as well as other androgens. Similarly, lower CYP19A1 activity caused by P228L variant of POR could lead to lower estrogen level. In addition, several steroid metabolism reactions are performed by CYP3A family of enzymes, especially, CYP3A4 in liver, and therefore, lower activities of CYP3A enzymes observed in our study by P228L variant of POR indicate a wider overall impact on multiple steroid metabolism pathways, affecting both androgen and estrogen production. Since POR is involved in large number of metabolic reactions catalyzed by cytochromes P450 enzymes in the endoplasmic reticulum, potentially, other xenobiotic and drug metabolizing reactions are also likely to be affected by the P228L variation in POR.

A single change in the amino acid sequence can affect protein stability and cause a significant reduction in POR activity. Molecular characterization of individual POR mutations is crucial to have a better understanding of the impact on the function of its redox partners.

## 4. Materials and Methods

### 4.1 Analysis of POR P228L allele frequencies

Allele Frequencies of POR P228L (DBSNP ID: rs17853284) were extracted from different databases: GnomAD_exome and GnomAD (Genome Aggregation Database), ALFA (Allele Frequency Aggregator), ExAC (Exome Aggregation Consortium), PAGE_STUDY (prenatal assessments of genomes and exomes), GO-ESP (Exome Variant Server), 1000 Genomes.

### 4.2 Conservation analysis

We performed ConSurf analysis to assess the evolutionary conservation of amino acids at position 228 in the structure of POR [24]. We aligned and collected 250 homologs of POR from the UniProt database using the Clustal Omega program. In the ConSurf server, we set as reference human POR amino acid sequence from UniProt database (ID #P16435) and crystal structure from PDB database (ID # 5FA6 chain A).

### 4.3 In silico predictions of POR P228L stability

We predicted the impact of the variant P228L using a set of sequence-based in silico tools, PredictSNP2, CADD, DANN, FTHMM, FunSeq2, GWAVA, and Meta-SNP [36, 37] using the human POR reference sequence (NCBI: NP_000932.3). Additionally, we used DynaMut-2.0 [38] tool to evaluate how the single amino acid change in POR could affect its structure, function, and interactions, based on its crystal structure data (PDB ID # 5FA6 chain A).

### 4.4 Protein structure analysis of POR variants

Three-dimensional structural models of POR (NP_000932) proteins were obtained from the protein structure database (www.rcsb.org). We used the structures of the human POR (PDB # 5FA6) to analyze the location of amino acids described in this report [8]. Structure models were drawn using Pymol (www.pymol.org).

### 4.5 Expression of POR in E. coli

WT or P228L POR variant cDNAs with a His-tag were cloned in a pET22b and transformed into Escherichia coli BL21(DE3), single colonies were selected by growing in LB media with 100 μg/mL carbenicillin. The large-scale expression was done by autoinduction system [39], growing the selected colonies in terrific broth supplemented with glucose 0.05 %, lactose 0.2 %, succinate 20 mM, NaSO_4_ 5 mM, NH_4_Cl 50 mM, MgSO_4_ 2 mM, 0.05 mg/ml riboflavin, and 100 μg/ml carbenicillin at 37 °C to an optical density (OD at 600nm) of 0.6 and then the temperature was reduced to 24 °C and cultures were grown for further 16h with constant shaking. The bacterial cells were collected by centrifugation, washed with PBS, and suspended in 50 mM potassium phosphate (pH 7.6) containing 250 mM sucrose, 0.5 mM EDTA, 0.2 mg/ml lysozyme, 1 mM PMSF and 20 U/ml Endonuclease for 1 h with slow stirring to generate spheroplasts. The spheroplasts were pelleted by centrifugation at 5000×g for 20 min; and suspended in 50 mM potassium phosphate (pH 7.8) containing 6 mM MgOAc, 0.1 mM DTT, 20% (v/v) glycerol, and 1 mM PMSF; and disrupted by sonication. A clear lysate devoid of cellular debris was obtained by centrifugation at 12,000×g for 10 min, and then the membranes were collected by centrifugation at 100,000×g for 90 min at 4 °C. Membranes containing over-expressed POR WT or P228Lvariant were stored at −80 ◦C.

### 4.6 Purification of recombinant human POR from isolated E. coli membranes

All steps were carried out at 4 °C. His-tagged proteins were solubilized at a concentration of 0.25 g of membrane per mL of 50 mM potassium phosphate, pH 7.4, 10% (v/v) glycerol, 1% Triton X-100. The mixture was gently stirred for 16 h and then centrifuged at 12,000 ×g for 15 min and the supernatant was used for purification by ion-metal affinity chromatography (IMAC). The supernatant was diluted with buffer A to a final concentration of 50 mM potassium phosphate, pH 7.4, 30 mM imidazole, 0.1 % Triton, 150 mM NaCl and 10 % glycerol. This mixture was loaded in 4 mL His60 Ni Superflow™ Resin and the impurities were washed with buffer A with an increasing concentration of imidazole to 60 mM. The His-tagged proteins were eluted with the same buffer with increasing concentrations of imidazole up to 500 mM and the presence of POR was confirmed by Western blot. Purified samples were concentrated, and the elution buffer was exchanged with 50 mM Potassium Phosphate, pH 7.4, 10% (v/v) glycerol, 0.1% Triton X-100 using Amicon® Ultra Centrifugal Filters 20,000 MWCO. Protein concentration was measured by Bradford [4, 15, 34, 40].

### 4.7 Flavin content

Protein samples were treated with 2 M urea to release the Flavin molecules from the protein structure. Then, precipitated proteins were removed by centrifugation at 13 000 rpm for 10 minutes. The fluorescence of released FMN and FAD was measured at pH 7.7 and 2.6 respectively (excitation at 450 nm, emission at 535 nm) to determine the Flavin content [41].

### 4.8 Fast proteolysis assay (FASTpp)

The FASTpp was performed as previously described with some small modifications [29, 42]. Membranes were solubilized using 1 % Triton X-100 and centrifuge at 12000 g for 40 minutes. The supernatant was used for the next steps. The reaction mixture contained 0.4 mg/ml of solubilized membranes expressing WT or P228L POR, 0.05 mg/ml of Thermolysin (Sigma-Aldrich) in 10 mM CaCl_2_, 20 mM potassium phosphate buffer (pH 7.6) and digestion was performed in a Gradient Thermo block (Biometra) generating a gradient from 4°C to 70°C. After proteolysis samples were centrifuged at 12000 g to remove all the aggregated proteins. Samples were analyzed by Western Blot as described before [21]. We determined the inflection point of the melting curves, which is equal to the melting temperature (Tm), and a Boltzmann sigmoidal equation was fitted to the normalized fluorescent data. The data were analyzed with the GraphPad Prism program (GraphPadPrism v.3.00 for Windows, GraphPad Sofware, San Diego, CA, USA).

### 4.9 CPM fluorescence assay

CPM (diethylamino-3-(4-maleimidophenyl)-4-methylcoumarin) fluorescence assay was performed as described previously, with few modifications [43]. The working solution of CPM in DMSO (4 µM) was freshly made and all the reagents were kept at 4 ◦C till the beginning of the assay. 2.5 µM of CPM was mixed with 1 µM of POR WT or mutant, 50 mM of Tris-HCL buffer (pH 7.4), and increasing concentration of urea from 0.125 M to 1 M. The fluorescence signal was measured at an emission of 463 nm.

### 4.10 POR assays with cytochrome c and MTT

The assays were done in triplicates in 96-well format using a Spectramax M2e microplate reader (Molecular Devices, Sunnyvale, CA, USA). Each reaction well was composed of 50 nM POR in 50 mM Tris-HCl (pH 7.8), 150 mM NaCl, and cytochrome c concentration was varied from 2.5-60 μM. The reaction was started by the addition of 100 µM NADPH and monitored at 550 nm using the extinction coefficient (ε_550=_ 21.1 mM^-1^ cm^-1^) for 10 minutes. The reaction rates were extrapolated from the linear range of the kinetic traces and plots of rate versus cytochrome c concentration) were fitted with the Michaelis-Menton equation using PRISM (GraphPad, San Diego, CA) to determine Vmax and Km [21]. NADPH-dependent reduction of 3-(4,5-dimethylthiazol-2-yl)-2,5-diphenyltetrazolium (MTT) was measured as the rate of increase in absorbance at 610 nm using the extinction coefficient, ε_610_=11 mM^-1^ cm^-1^ (4). The assay mixture contained 50 nM POR in 50 mM Tris-HCl (pH 7.8), 150 mM NaCl, and MTT varied from 3.9-500 µM. The reaction was started by the addition of 100 µM NADPH [6, 15].

### 4.11 Assay of cytochrome P450 CYP3A4, CYP3A5, and CYP2C9 activity in reconstituted liposomes

The activity of CYP3A4, CYP3A5, or CYP2C9 in presence of WT or mutant POR was tested using the fluorogenic substrate BOMCC (Invitrogen Corp, Carlsbad, CA, United States). The purified CYP3A4, CYP3A5 or CYP2C9 (CYPEX, Dundee, Scotland, United Kingdom) were used to test the activities of the POR variants using 20 μM BOMCC as substrate. In vitro CYP3A5 assays were performed using a reconstituted liposome system consisting of pure WT/mutant POR, CYP3A5/CYP2C9 and cytochrome b_5_ at a ratio of 5:1:1. The final assay mixture consisted of 20 µM DLPC (1,2-Dilauroyl-snglycero-3-phosphocholine)/DLPGV (1,2-Dilauroyl-sn-glycero-3-phosphoglycerol) and proteins (100 nM POR: 20 nM CYP2C9: 20 nM b_5_), 3 mM MgCl2, 20 μM BOMCC in 50 mM Tris-HCl buffer pH 7.4 and the reaction volume was 100 µL. The CYP3A5 reaction was started by the addition of NADPH to 1mM final concentration, and fluorescence was measured on a Spectramax M2e plate reader (Molecular Devices, Sunnyvale, CA, United States) at an excitation wavelength of 415 nm and an emission wavelength of 460 nm for BOMCC [21, 31, 33].

### 4.12 Assay of cytochrome P450 CYP17A1 and CYP19A1

Activities of CYP17A1 and CYP19A1 were measure with WT or P228L variant of POR using methods described in detail in our previous publications [14, 18, 20, 21, 30, 40, 44, 45].

### 4.13 Statistical analysis of results

Data are shown as mean and standard errors of the mean (SEM) for each group of replicates.

## 5. Conclusions

Considering all available data from different functional assays, we suggest that rs17853284, the variant causing amino acid change P228L in POR is a disease-causing variant whose effect will be prominent when present together with another variant of POR on the second allele that has a severe impact on POR activities. A combination of rs17853284 with a severe mutation in POR (leading to a truncated protein, loss of flavins, and protein instability) has the potential to cause P450 oxidoreductase deficiency.

## Author Contributions

Conceptualization, A.V.P.; methodology, A.V.P., M.N. and M.N.R.V.; formal analysis, A.V.P. and M.N.R.V.; investigation, M.N. and M.N.R.V; resources, A.V.P.; writing—original draft preparation, M.N. and M.N.R.V.; writing—review and editing, A.V.P.; visualization, M.N. and M.N.R.V.; supervision, A.V.P.; project administration, A.V.P.; funding acquisition, M.N.R.V. and A.V.P. All authors have read and agreed to the published version of the manuscript.

## Funding

This research was funded by SWISS NATIONAL SCIENCE FOUNDATION, grant number 310030M_204518 to AVP. MNRV is funded by a SWISS GOVERNMENT EXCELLENCE SCHOLARSHIP (ESKAS) grant number 2020.0557.

## Notes

### Competing Interest Statement

The authors have declared no competing interest.

### Summary of Updates

Edited typographical errors, and updated title.

